# Nonsinusoidal oscillations underlie pathological phase-amplitude coupling in the motor cortex in Parkinson’s disease

**DOI:** 10.1101/049304

**Authors:** Scott R. Cole, Erik J. Peterson, Roemer van der Meij, Coralie de Hemptinne, Philip A. Starr, Bradley Voytek

**Affiliations:** Neurosciences Graduate Program, University of California, San Diego, 9500 Gilman Drive La Jolla, CA, 92093.; Department of Cognitive Science, University of California, San Diego, 9500 Gilman Drive La Jolla, CA, 92093.; Institute for Neural Computation, University of California, San Diego, 9500 Gilman Drive La Jolla, CA, 92093.; Kavli Institute for Brain and Mind, University of California, San Diego, 9500 Gilman Drive La Jolla, CA, 92093.; Department of Neurological Surgery, University of California, San Francisco, 505 Parnassus Ave, San Francisco, CA, 94143

**Keywords:** oscillation, phase-amplitude coupling, Parkinson’s disease, electrocorticography, motor cortex, beta, waveform, shape

## Abstract

Parkinson’s disease (PD) is associated with abnormal beta oscillations (13-30 Hz) in the basal ganglia and motor cortex (M1). Recent reports show that M1 beta-high gamma (50-200 Hz) phase-amplitude coupling (PAC) is exaggerated in PD and is reduced following acute deep brain stimulation (DBS). Here we analyze invasive M1 electrocorticography recordings in PD patients on and off DBS, and in isolated cervical dystonia patients, and show that M1 beta oscillations are nonsinusoidal, having sharp and asymmetric features. These sharp oscillatory beta features underlie the previously reported PAC, providing an alternative to the standard interpretation of PAC as an interaction between two distinct frequency components. Specifically, the ratio between peak and trough sharpness is nearly perfectly correlated with beta-high gamma PAC (*r* = 0.96) and predicts PD-related motor deficit. Using a simulation of the local field potential, we demonstrate that sharp oscillatory waves can arise from synchronous synaptic activity. We propose that exaggerated beta-high gamma PAC may actually reflect such synchronous synaptic activity, manifesting as sharp beta oscillations that are “smoothed out” with DBS. These results support the “desynchronization” hypothesis of DBS wherein DBS counteracts pathological synchronization throughout the basal ganglia-thalamocortical loop. We argue that PAC can be influenced by more than one mechanism. In this case synaptic synchrony, rather than the often assumed spike-field coherence, may underlie exaggerated PAC. These often overlooked temporal features of the oscillatory waveform carry critical physiological information about neural processes and dynamics that may lead to better understanding of underlying neuropathology.

## Introduction

Parkinson’s disease (PD) is characterized by neuronal degeneration in multiple systems, including midbrain dopaminergic neurons. Though beta (13-30 Hz) oscillations are a normal feature of the basal ganglia-thalamocortical loop, PD is associated with excessive neuronal synchronization in the beta band (1, 2). Despite an established relationship between beta band neuronal synchronization and PD, the physiological mechanism causing motor dysfunction has been unclear. Excessive phase-amplitude coupling (PAC) between beta phase and high gamma amplitude (50-200 Hz) may offer an explanation (3–5). PAC between distant neural populations has been linked to enhanced neural information flow (6–8), long-term potentiation (9), and improved behavioral performance (10). However PAC strength is greater in M1 of PD patients compared to patients with isolated cervical dystonia or epilepsy (3), leading to the hypothesis that beyond its facilitative role, PAC may play a role in neural pathology (3, 4, 11).

Analyses of PAC implicitly presuppose two separate, interacting physiological processes: a low-frequency component associated with an oscillation in the synaptic currents and a separate high-frequency component associated with local spiking activity. The degree of interaction between these two signals is then quantified using a single PAC metric, often with the assumption that low frequency oscillatory phase organizes neuronal cell assembly spiking (3, 4, 8, 12–14).

However the spectral features indicative of PAC can also arise from different temporal features in data, such as sharp temporal deflections in the local field potential (LFP) (15, 16) produced by synchronous synaptic activity (17, 18). That is, rather than PAC reflecting two distinct, interacting processes, one asymmetric, nonsinusoidal oscillation will also manifest PAC. In order to adjudicate between these two possible mechanistic interpretations of pathological PAC in PD, we characterize the shape of oscillatory waveforms in 23 PD patients on and off DBS (data from (4)), as well as from 9 patients with isolated cervical dystonia without arm involvement (data from (3)).

We report that beta-high gamma PAC in PD does not primarily arise from the coupling between a beta oscillation and a high-frequency asynchronous signal. Instead, M1 PAC is almost fully captured by changes in the shape of the beta waveform. We further show that the beta waveform is “smoothed out” with DBS treatment, differs between PD and isolated cervical dystonia patients, and predicts PD patient rigidity. Further, we utilize a simplified simulation of field potential data to show that altered waveform shape can arise via changes in the synchronization of synaptic currents, consistent with empirical reports (17, 18).

## Results

We characterize the shape of beta waveforms by measuring the sharpness of the peaks and troughs (see SI Methods). Specifically, we quantify the symmetry of these waveforms by defining the extrema sharpness ratio (ESR). ESR is the ratio between the average peak sharpness and the average trough sharpness over all peaks and troughs in the 30-second recording. This ratio is fixed to be strictly greater than 1 such that an oscillation with high peak-trough asymmetry will have a value higher than 1 regardless of whether the peak is sharper than the trough or vice versa (see SI Methods).

In intracranial electrocorticography recordings from arm area of M1, we show that waveform shape differs between PD and isolated cervical dystonia patients. We further show that DBS treatment of PD alters waveform shape. Using the covariation between two aspects of waveform shape, extrema sharpness and rise/decay steepness, we characterize the stereotypical waveform of M1 beta oscillations as a type of sawtooth. We compare measures of ESR and PAC across subjects to show that these measures are highly correlated and reflect the same aspects of the electrophysiological signal despite analyzing different domains (time and frequency, respectively). We summarize by presenting a computational model in which increased synchrony of synaptic events biases both oscillatory waveform shape and PAC.

### Waveform shape of M1 beta oscillation changes in PD

We apply the ESR measure to characterize the waveform shape of M1 beta oscillations in two patient groups: PD and cervical dystonia. ESR is greater in PD compared to cervical dystonia (Fig. 1A, Mann-Whitney *U* test, *U*_30_ = 48 *ρ* = 0.021), meaning that M1 beta oscillations in PD are more asymmetric in regards to the sharpness of the peaks and troughs. Additionally, DBS treatment decreases ESR in PD patients (Fig. 1B-D, paired *t*-test, *t*_22_ = 2.6, *ρ* = 0.016), so that the ESR is closer to that in the cervical dystonia patients. Furthermore, PD patients’ clinical rigidity scores pre-DBS positively correlate with ESR (Fig. 1E, Spearman *r* = 0.59; *n* = 23; *ρ* = 0.006). Similar relationships between PD and waveform shape were observed for a second measure of waveform shape, the rise-decay steepness ratio (see SI Methods, Fig. S1A-C). Thus, there is a characteristic difference in the shape of beta oscillations in untreated PD patients compared to cervical dystonia patients or PD patients treated with DBS.

**Fig. 1.**
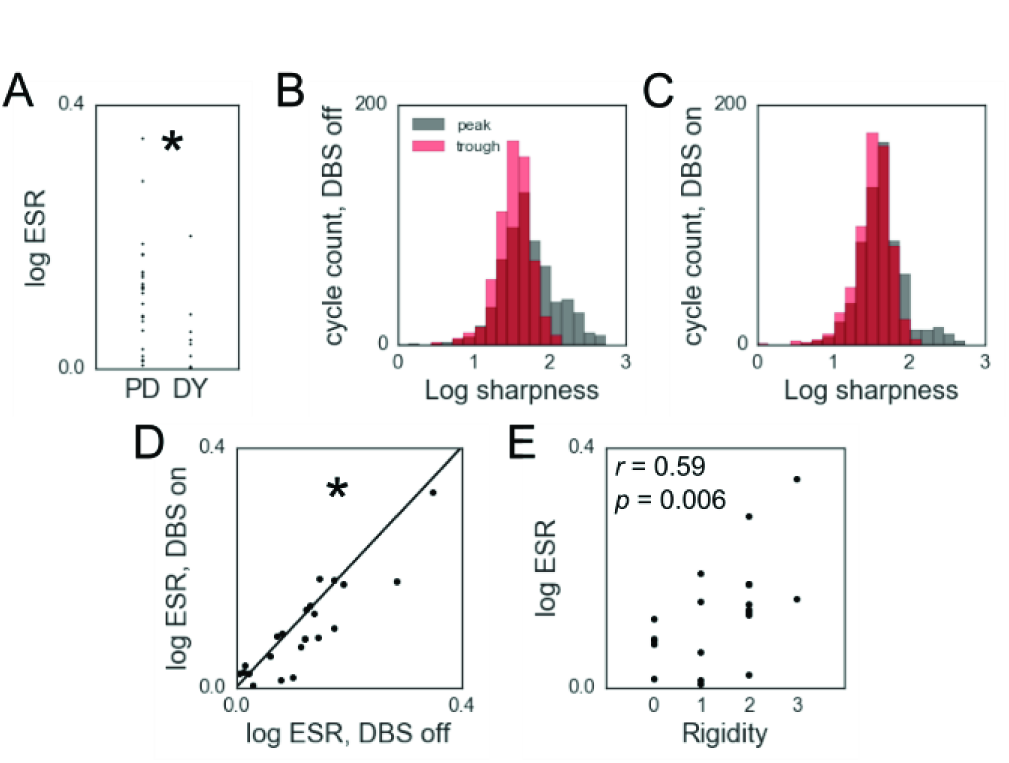
Waveform shape of M1 beta oscillation changes in PD. (A) ESR is higher in PD patients compared to dystonia patients. (B,C) Distributions of peak and trough sharpness before (B) and during (C) DBS for an exemplar patient. Note that there is greater overlap between these distributions with DBS on. (D) ESR decreases in PD patients with DBS application. Diagonal line represents unity. (E) Clinical rigidity scores are positively correlated with ESR in PD patients pre-DBS. Each dot represents 1 patient in panels A, D, and E. * indicates p < 0.05.

It is important to note that ESR can be changed by increases and/or decreases in the sharpness of one or both extrema. We find that M1 beta oscillation extrema are sharper in PD compared to cervical dystonia (Fig. S2A; Mann-Whitney *U* test, *U*_62_ = 171, *ρ* = 0.0003). Additionally, DBS trends toward flattening sharp extrema as opposed to sharpening flat extrema (Fig. S2B; paired *t*-test, *t*_45_ = 1.8, *ρ* = 0.071).

### M1 beta waveform is a characteristic sawtooth

The sharpness ratio between peaks and troughs was compared to the steepness ratio between rises and decays, across subjects. We report a strong correlation between these measures (Fig. 2A, *r* = 0.85, *ρ* < 10^−23^). That is, patients whose peaks are sharper relative to their troughs also have rise phases that are relatively steeper than their decay phases (Fig. 2B, top right). In contrast, patients whose troughs are sharper than their peaks have decay phases that are steeper than their rise phases (Fig. 2B, bottom left). Furthermore, these waveforms are the kind expected if the LFP was heavily influenced by synchronous synaptic potentials (see Fig. 4A, simulated data), *i.e.*, it is similar to the rise-and-decay profile of individual postsynaptic potentials. This waveform shape is consistent with the dark gray, but not the light gray, sawtooth shapes in Fig. 2B.

**Fig. 2.**
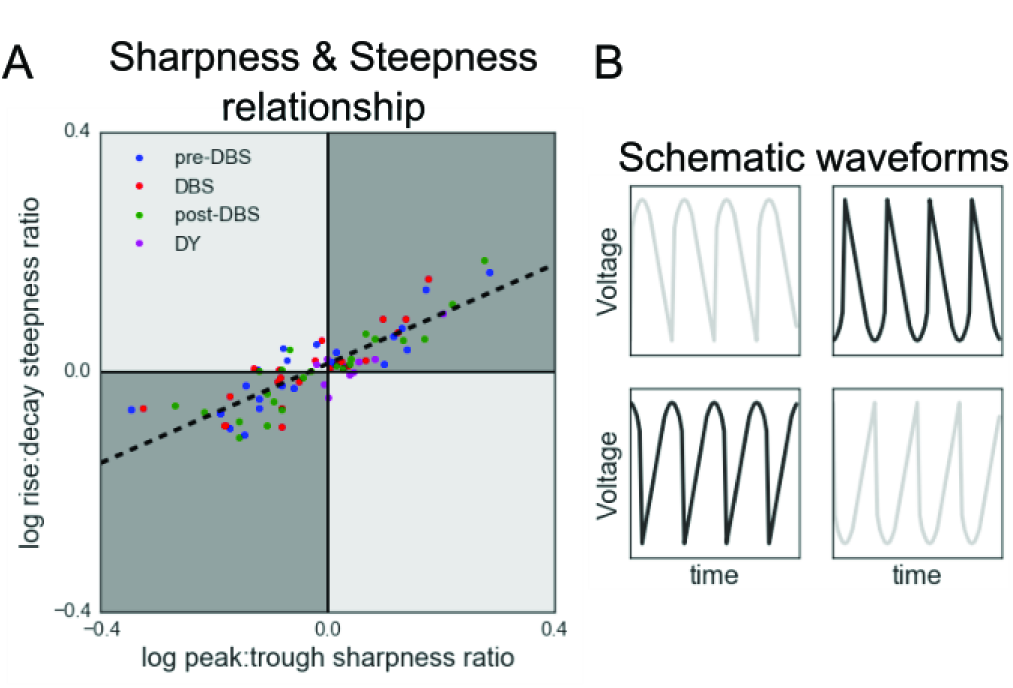
M1 beta oscillations have a characteristic sawtooth shape. (A) Positive correlation between the relative sharpness of a patient’s oscillatory peaks and the relative steepness of a patient’s voltage rises. Each marker represents one 30-second recording from a PD or dystonia patient (see legend). (B) Schematic voltage traces corresponding to each quadrant of (A). M1 beta falls in the gray quadrants (quadrants I and III) of this two-dimensional space.

**Fig. 3.**
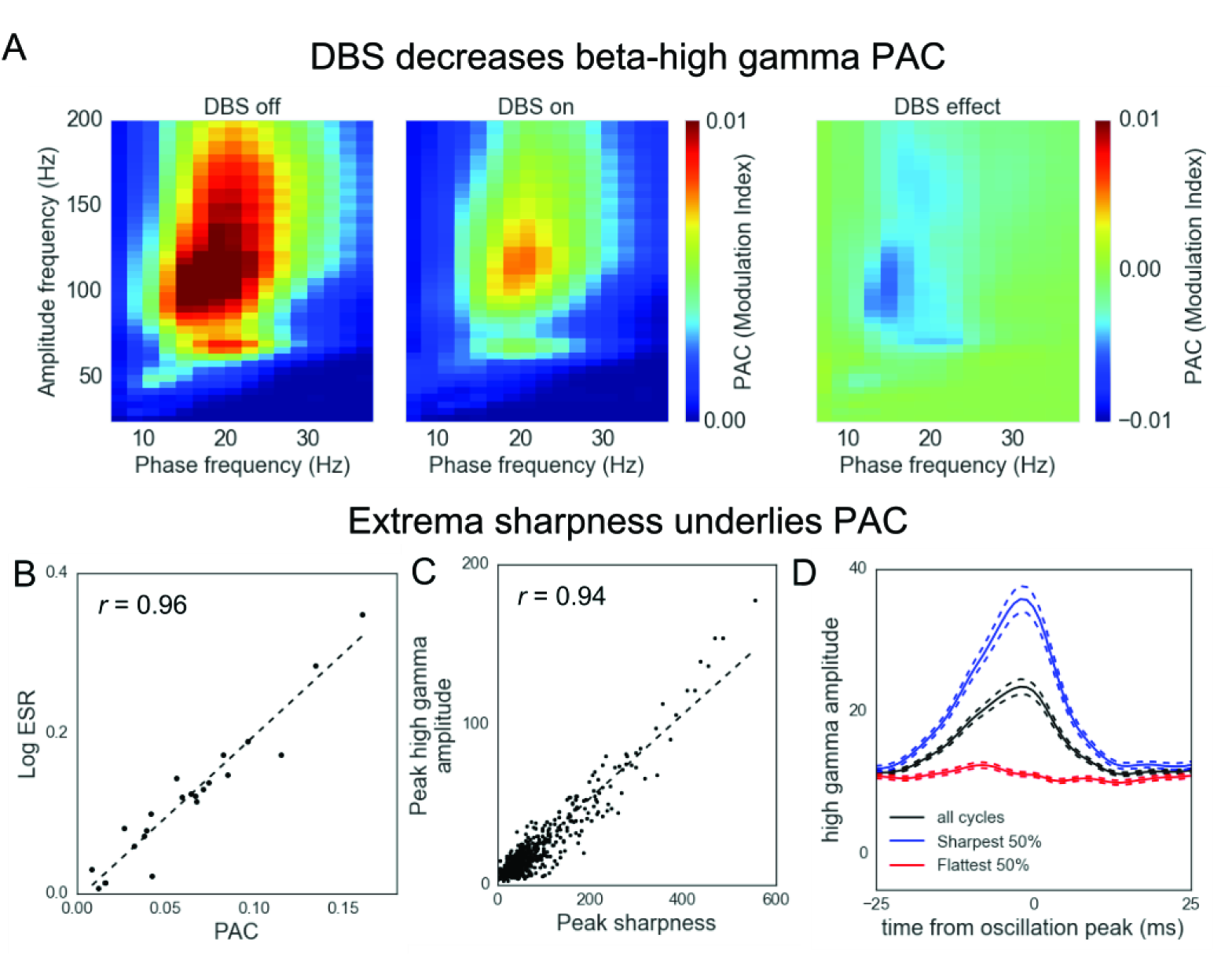
Extrema sharpness of beta oscillations underlies PAC. (A) Example patient showing a decrease in beta-high gamma PAC with DBS application, as described previously (4). (B) Strong correlation between the magnitude of beta-high gamma PAC and ESR. A value of 0 represents equal sharpness between peaks and troughs. Each dot represents one patient. (C) Positive correlation between the sharpness of a peak and the high gamma amplitude at that time. Each dot represents one peak in the 30-second recording of the exemplar PD patient in panel A, pre-DBS. (D) High gamma amplitude time-locked to oscillatory peaks in the 30-second recording of the same exemplar PD patient pre-DBS. Maximal high gamma was observed at the peak of the oscillation (*i.e.*, PAC). This effect was observed when analyzing all oscillations (black). However, it was strongest when restricting analysis to the 50% of cycles with sharpest peaks (blue), and was not observed for the 50% of cycles with flattest peaks (red). Solid lines denote mean and dashed lines denote SEM.

**Fig4.**
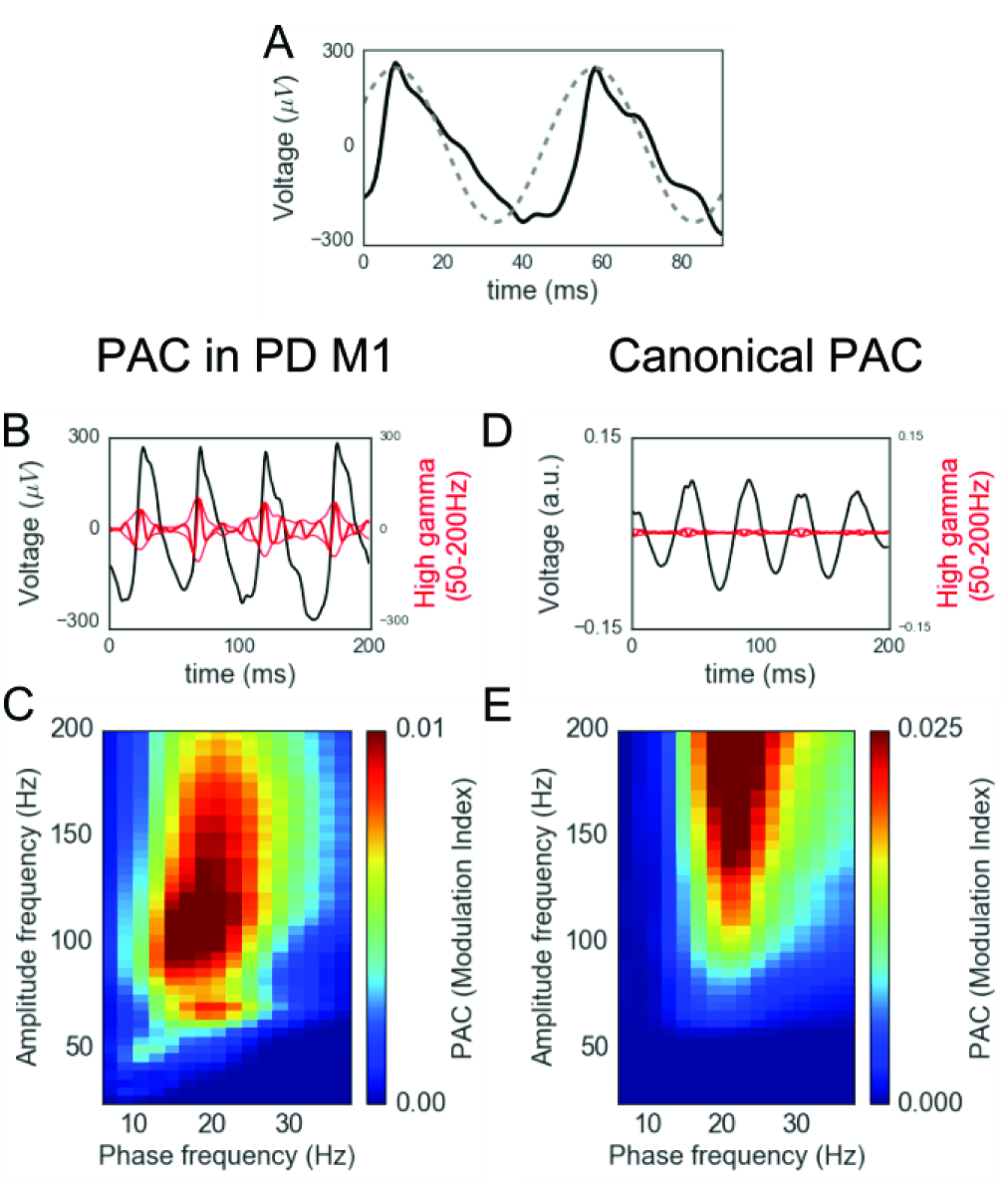
Nonsinusoidal waveforms produce PAC. (A) Exemplar unfiltered beta oscillation recorded from a PD patient pre-DBS (black), contrasted with a sinusoid of comparable frequency (gray). (B) Overlay of an exemplar raw voltage recording from a PD patient pre-DBS (black) and its high gamma component (red). The thick red line is the black trace band-pass filtered in the high gamma range (50-200 Hz), and the thin red lines show the analytic amplitude of this component. Note the black and red traces are on the same voltage scale, showing that high gamma is much larger in amplitude than normally associated with high gamma that arises from total population spike rate (18, 19). (C) Phase-amplitude coupling comodulogram of the 30-second recording from the PD patient pre-DBS shown partially in panels A & B. (D) Simulated canonical beta-high gamma phase-amplitude coupling in which the high gamma oscillation has greater amplitude around the peak of the beta oscillation. (E) Comodulogram of the simulated 30-second signal shown partially in panel D. Note its similarity to the comodulogram in panel C despite the different appearance between the signals in the time domain (panels B, D).

### Waveform shape of beta oscillations underlies PAC

M1 beta-high gamma PAC is increased in PD patients relative to epilepsy and cervical dystonia patients (3) and decreases with DBS application (Fig. 3A) (4). Additionally, we note in untreated PD patients that high gamma amplitude is specifically coupled to the peaks and troughs of the beta oscillations, as opposed to non-extrema phases of the beta cycle (Fig. S3).

Because sharp transients in the local field potential are known to yield PAC (15, 16), we correlated ESR and PAC across subjects. We find that ESR is strongly correlated with PAC across PD patients (Fig. 3B; *r* = 0.96; *ρ* < 10^−12^). This is because beta sharpness and apparent high gamma amplitude are related to the same features of the electrophysiological signal, shown by the strong correlation between peak sharpness and its instantaneous high gamma amplitude in an example patient (Fig. 3C, pre DBS; *r* = 0.94; *ρ* < 10^−291^). This holds true for both peaks and troughs of all PD patients (Fig. S4A,B). Furthermore, high gamma amplitude is time-locked to sharp extrema, but not flat extrema, shown in the same example patient (Fig. 3D; Fig. S5 for all patients). Similar results were observed when comparing PAC to the rise-decay steepness ratio (Fig. S1D,E and Fig. S4C,D). While the analysis in Fig. 3B was calculated using pre-DBS recordings, similar relationships were observed in recordings during and after DBS as well as in patients with cervical dystonia (Fig. S6).

Furthermore, ESR was calculated on data that was low-pass filtered at 50 Hz, removing the high gamma component used to calculate PAC. Despite negligible high gamma amplitude being present in the filtered signal on which ESR was calculated, ESR is still correlated with PAC (*r* = 0.69, *ρ* < 0.001), showing that this correlation is not an artifact of the presence of high gamma (Fig. S7).

Because ESR is calculated using the raw electrophysiological signal of each extremum and 2 samples (sampled at 1 kHz) around it, 90% of the variance is captured by only using about 12% of the data. However, ESR is not the only dimension of shape that correlates with M1 PAC, as other features of the waveform will ultimately determine the results of sinusoidal decomposition (such as RDSR). ESR and RDSR both individually correlate with PAC after holding the other metric constant (partial correlations, ESR: *r* = 0.71, *ρ* < 0.001; RDSR: *r* = 0.46, *ρ* = 0.028) and together explain 95% of the variance in PAC of PD patients pre-DBS.

In summary, the time-domain shape of a low frequency oscillatory waveform can manifest as PAC (Fig. 4A-C). While high gamma amplitude has been shown to correlate with local population spiking activity (18, 19), the magnitude of these “true” high gamma changes are low, on the order of a few microvolts (Fig. 4D,E). In contrast, apparent high gamma resulting from sharp time-domain deflections is an order of magnitude stronger–nearly 100 microvolts in some cases (Fig. 4B).

### Relationship between beta power and waveform shape

The power in a frequency band is a commonly reported measure that may change with disease state and may covary with other metrics. We quantified the effect of DBS on beta and high gamma power. Across all patients, there is a trend towards DBS decreasing power in both beta (Fig. S8A; paired t-test; *ρ* = 0.11) and high gamma (Fig. S8B; paired t-test, *ρ* = 0.076) bands. Furthermore, the beta power change with DBS is positively correlated with the ESR change across patients (Fig. S8C; *r* = 0.57; *ρ* = 0.004). As expected, a positive correlation was also observed between beta power and PAC (Fig. S8D; *r* = 0.48; *ρ* = 0.022). However, although beta power positively correlates with both ESR and PAC, ESR is still correlated with PAC after holding beta power constant (partial correlation, ESR: *r* = 0.89, *ρ* < ^−7^).

### Sharp oscillations are generated by synchronous synaptic currents in a computational model

We simulated LFPs in attempt to relate our measure of oscillation shape to the underlying physiology. Simulated neurons randomly generated synaptic events following a time-varying rate (see SI Methods). By adjusting the mapping between simulated beta phase and synaptic event rate, we manipulated the rhythmic synchrony of synaptic events (see SI Methods). Synchronous synaptic events yield sharper local field potentials compared to non-synchronous currents (Fig. 5A). When observing several periods of fluctuations in synaptic activity, increased synaptic synchrony at a beta rhythm causes sharper oscillatory beta waveforms in the LFP (Fig. 5B). High synaptic synchrony yields both increased ESR (Fig. 5C) and beta-high gamma PAC (Fig. 5D). Therefore, high synaptic synchrony produces simulated M1 field potential recordings similar to those of PD patients without treatment, and low synaptic synchrony produces local field potentials more similar to cervical dystonia patients or PD patients with DBS treatment.

**Fig. 5.**
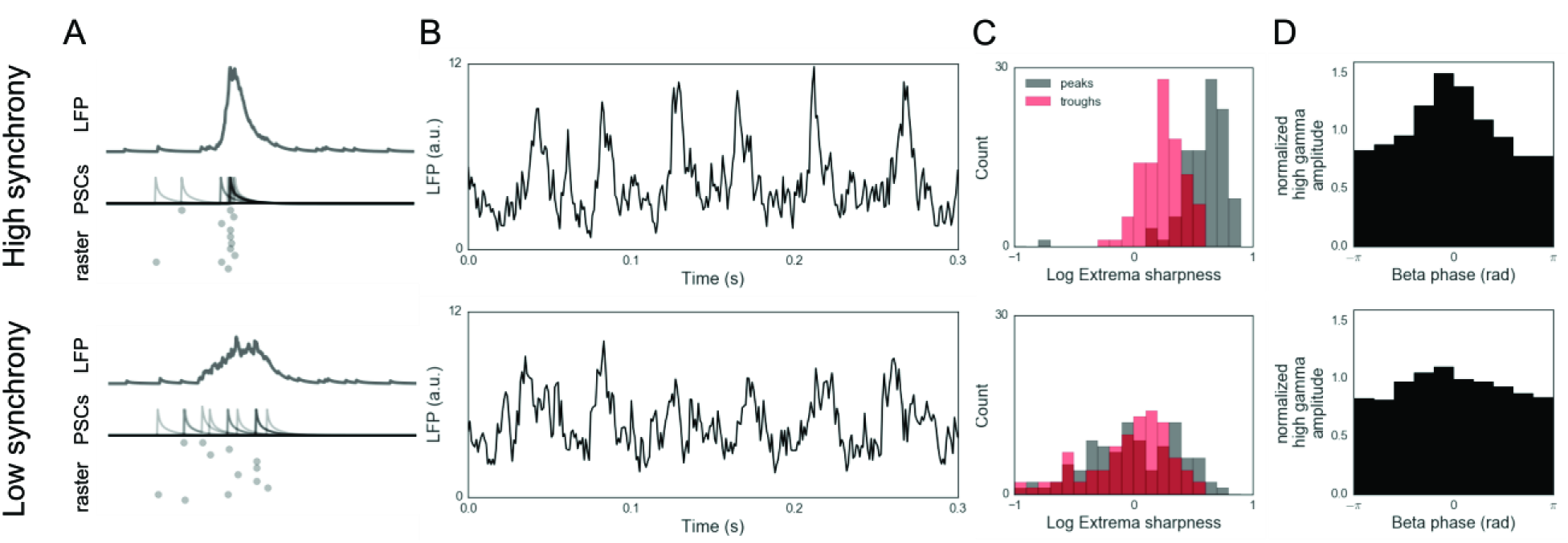
Synchrony of synaptic currents affects oscillation shape in a simulated neuron population. (A) Top: In a regime of high synaptic synchrony across cells in the raster plot, postsynaptic currents (PSCs) summate in order to generate a sharp deflection in the local field potential (LFP). Bottom: This deflection is flatter when synaptic currents are less synchronous. (B) Simulated LFPs from neural populations that have rhythmic activity at a beta frequency. Highly synchronous activity yields sharper beta oscillations (top) compared to lower synchrony (bottom). (C) Distributions of peak and trough sharpness for the simulated LFPs in panel B. Note the greater overlap between extrema sharpness distributions in the lower synchrony condition. (D) Coupling between beta phase and high gamma amplitude for the simulated LFPs in panel B. Note that coupling is greater in the simulated LFP with high synaptic synchrony.

## Discussion

It has been hypothesized that the key mechanism by which DBS reduces PD-related movement symptoms is by the decorrelation of the excessively synchronized neural activity in the basal ganglia (11, 20–22), consistent with the previously reported PAC decrease with DBS treatment (4, 5). The present work offers new insight into the biophysical interpretation of PAC, as arising from a decrease in the synchrony of synaptic currents in M1, which are likely a mix of both afferent and local events. Similar to previous interpretations of therapeutic reduction in PAC (4, 5), decreased synaptic synchrony would give top-down, executive input from the prefrontal cortex increased efficacy in biasing M1 spiking. As such, DBS acts to “free up” communication channels into M1, channels that were previously dominated by the pathological oversynchronization of synaptic inputs. Given how potent synchronous activity can be in driving downstream population spiking (23–25), we suggest that excessive synchrony in presynaptic inputs to M1 may be key to understanding the biophysical measurements related to excessive cortical PAC. Of note, we have argued that current DBS treatment could potentially be improved by applying feedback from motor cortical activity (4). Detection of a sharp waveform in the motor cortical field potential as a trigger for adaptive stimulation, would be both computationally simpler than calculating PAC and a more direct measure of the pathological biomarker.

Our work demonstrates that PAC, as an analytic measure, is sensitive to two different but related physiological phenomenon. First, beta-high gamma PAC has been interpreted as a true coupling between a low frequency oscillation and a distinct physiological process that generates amplitude at high frequencies (local spiking). Here we show that PAC may also be generated from synaptic synchrony that generates a sharp low-frequency oscillation. From this perspective, it is clear that PAC as an analytic measure is not specific to one unique physiological phenomenon. This is critical, as PAC has been hypothesized to represent effective neural communications mediated by cross-frequency coupling between brain regions (8, 11, 12, 26) and is usually interpreted within the two-process framework, rather than the one-process alternative offered here (3, 4, 8, 12–14). In the past, increases in PAC have been associated with improved multi-item working memory (27), learning (10), attention (28), decision making (29), and cognitive control (8, 30). We have shown that excessive PAC in PD is generated by a single slow oscillatory waveform of variable shape, and offer the possibility that previous work showing PAC changes during normal cognition might also be driven by changes in synaptic input structure as well.

We hypothesize that the shape of neural oscillations carries information about the underlying neural activity. For example, the asymmetry of the hippocampal theta oscillation and cortical slow oscillations match the asymmetry of the firing rate histograms in their respective areas (31, 32). Because of the noticeable asymmetry, strategies have been developed to aid in the analysis of nonsinusoidal oscillations. For instance, theta oscillatory phase has been computed using features of the raw voltage (32–34). Additionally, empirical mode decomposition has been used to extract rhythmic components for phase-amplitude coupling instead of a Fourier decomposition (35). However, until now, techniques that characterize the shape of oscillatory waveforms are lacking.

Insight into the meaning of an oscillatory waveform is aided by recognizing that the LFP is predominantly composed of the spatiotemporal summation of synaptic potentials (36–40). Because synaptic potentials are both excitatory and inhibitory, the significance of an extracellular voltage deflection is not obvious, but nonetheless reflects fluctuations in the relative excitability of the local population. For example, a sharp decrease in the extracellular voltage could indicate synchronous activation of presynaptic glutamatergic projections to the recorded region. Alternatively, this voltage change could also be generated by a sudden cessation of local inhibition. Both of these changes, and other possibilities, could be supported or rejected by experiments with simultaneous measurement of additional signals, such as local spiking or presynaptic activity.

Oscillatory activity is abundant in electrophysiological recordings across many species, brain regions, and spatial scales (41–44), with canonical frequency bands (delta, theta, alpha, beta, gamma) often said to associate with (or even drive) specific behaviors and functions. Thus, the prospect that nonsinusoidal features of those oscillations carry critical physiological information provides a new framework for linking physiology with network dynamics and processes. For example, two studies have shown that the asymmetry of hippocampal theta oscillations changes with the type of movement in mice (45, 46).

In summary, we have shown that the shape of beta oscillations in field potentials measured from the arm-related area of M1 is changed by DBS and differs between PD patients and patients with isolated cervical dystonia and normal arm function. Further, ESR (and presumably, therefore, synaptic input synchrony) is positively correlated with a clinical measure of rigidity in PD. Using a computational model, we showed that the beta waveform shape is consistent with highly synchronous synaptic inputs and underlies the previously reported beta-high gamma PAC. We argue that both spectral and temporal analyses may be necessary to extract, and interpret, information in electrophysiological signals.

## Methods

Methods for collection of this data were previously reported (3, 4) and further details are available in the SI Methods. Briefly, intracranial recordings were collected from 23 PD patients with DBS on and off and 9 dystonia patients. PD patients were selected to have mild to moderate bradykinesia without a prominent tremor. A six-contact electrode strip was placed on the cortical surface through the burr hole used for DBS. A bipolar configuration was used for referencing. The electrode over M1 was localized with computed tomography and evoked somatosensory potentials. Data were downsampled to 1 kHz.

Prior to characterizing oscillation shape, peaks and troughs were identified in the raw time series. Sharpness of each extrema was estimated by averaging the voltage difference between the extrema and the samples 5ms before and after the extrema. Extrema sharpness ratio (ESR) was calculated as the ratio between the average peak and trough sharpness in the recording, and the ratio was fixed to be greater than 1. Rise and decay steepness were quantified as the steepest voltage change in 1 sample (millisecond) between each peak and trough. Rise-decay steepness ratio (RDSR) was the ratio between the average rise steepness and decay steepness throughout a recording.

Phase-amplitude coupling was calculated using the normalized modulation index metric (47). Oscillatory phase was estimated by the angle of the Hilbert transform of the signal bandpass filtered in the beta range (13-30Hz). Amplitude was similarly calculated by the magnitude of the Hilbert transform of the high gamma (50-200Hz) band-pass filtered signal.

Unless indicated otherwise, all correlations are Pearson and all tests are two-tailed.

## Author contributions

S.R.C. performed the research, S.R.C., E.J.P., R.v.d.M. and B.V. designed the research, C.d.H. and P.A.S and designed and performed the original PD study; all authors wrote the paper.

## Acknowledgements

We thank T. Donoghue, R. Gao, T. Noto, B. Postle, and T. Tran for invaluable discussion and comments. S.R.C. is supported by the National Science Foundation Graduate Research Fellowship Program. B.V. is supported by the University of California, San Diego, Qualcomm Institute, California Institute for Telecommunications and Information Technology, Strategic Research Opportunities Program, and a Sloan Research Fellowship. B.V. and R.v.d.M. are further supported by US National Institutes of Health grant NIH MH095984 to B.R. Postle. Data collection by P.A.S. and C.d.H. was supported by a grant from the Michael J. Fox foundation and by US National Institutes of Health grant R01 NS069779.

